# Genomic resources for comparative analyses of obligate avian brood parasitism

**DOI:** 10.1101/2025.08.12.669747

**Authors:** Rachel A. Carroll, Edward S. Ricemeyer, LaDeana W. Hillier, Jeffrey M. DaCosta, Ekaterina Osipova, Sara Smith, Gabriel A. Jamie, Juan G. Martinez, Mercedes Molina-Morales, Tomas Marques-Bonet, Joseph D. Manthey, Diana Haddad, Matthew J. Fuxjager, Kathleen S. Lynch, Jonathan M.D. Wood, Erich Jarvis, Patrick Masterson, Francoise Thibaud-Nissen, Mark Hauber, Claire N Spottiswoode, Timothy B. Sackton, Christopher N Balakrishnan, Michael D. Sorenson, Wesley C. Warren

## Abstract

Examples of convergent evolution, wherein distantly related organisms evolve similar traits, including behaviors, underscore the adaptive power of natural selection. In birds, obligate brood parasitism, and the associated loss of parental care behaviors, has evolved independently in seven different lineages, though little is known about the genetic basis of the complex suite of traits associated with this rare life history strategy. We generated genome assemblies for ten brood parasitic species plus eight species representatives of their parental/nesting outgroups. This includes nine long-read chromosome-level assemblies, with scaffold N50 sizes ranging from 38.1 to 72.6 MB, and gene representation completeness measures >97%. Leveraging this new catalog of avian genomes, we constructed clade-level alignments that reveal variation in chromosomal synteny, provide first-time or improved annotations of protein-coding and non-coding genes, and define cross-species ortholog reference sets. We also refine estimates for the timing of the seven independent origins of brood parasitism, ranging from recent events such as 1.6 to 4.5 million years ago in *Molothrus* cowbirds to much earlier origins over 30 million years ago in two of the three cuckoo lineages. These genomic resources lay the foundation for investigating the genetic and genomic underpinnings of brood parasitism, including the loss of parental care, shifts in mating systems, perhaps resulting in heightened sperm competition, elevated annual fecundity, improved spatial cognition related to nest-finding, and the diverse adaptations shaped by intense coevolution with host species.

## Introduction

About 100 avian species, or 1% of all taxa, are obligate brood parasites, reproducing only by laying their eggs in the nests of other bird species and thereby exploiting the parental care behavior of their “hosts” (Davies 2000). Obligate brood parasitism has independent origins in seven different lineages across four divergent avian orders (Payne 1977, Sorenson and Payne 2002), and so provides a compelling opportunity for comparative genomic analysis. Shared among these lineages is the loss of all parental care behaviors, including nest-building, incubation, and the brooding, provisioning, and defending own offspring.

Comparative genomic analyses of adaptation have often considered the convergent evolution of specific traits, such as novel sensory traits (Parker, Tsagkogeorga et al. 2013) (Sadanandan, Ko et al. 2023, Zou, Huang et al. 2024), adoption of new diets (Hu, Wu et al. 2017, Toda, Ko et al. 2021, Eliason, Mellenthin et al. 2023), or invasion of extreme environments (Foote, Liu et al. 2015, Cole, Zhou et al. 2022) (Lyu, Zhou et al. 2022). In other cases, convergent adaptation involves the loss of pre-existing traits such as flight (Sackton, Grayson et al. 2019) or functional vision (Moran, Richards et al. 2023). In comparison, the loss of parental care in parasitic birds represents a major behavioral and life history transition with multiple axes of potential convergence. In addition to the loss of parental behaviors, a substantial increase in female fecundity is another trait likely shared across all seven parasitic lineages is (Payne 1974, Payne 1976), reflecting a reallocation of time and energy from parental care to egg production (Payne 1977). Other axes of possible convergence are not as well characterized, but may include altered sexual selection associated with loss of parental care (Feeney and Riehl 2019), better spatial ability associated with finding and tracking the status of host nests (Sherry, Forbes et al. 1993, Reboreda, Clayton et al. 1996, Nair-Roberts, Erichsen et al. 2006, Guigueno, Snow et al. 2014) and accelerated pre- and/or post-hatch development (Birkhead, Hemmings et al. 2011) and novelties in behavioral imprinting and song learning (Payne, Payne et al. 1998, Louder, Balakrishnan et al. 2019). Many brood parasitic birds and their hosts also serve as important models for coevolutionary arms-races and their reciprocal interactions have produced a spectacular diversity of behavioral, morphological and physiological adaptations and counter-adaptations e.g. (Langmore, Hunt et al. 2003) (Yang, Liang et al. 2010, Spottiswoode and Stevens 2011) (Langmore, Stevens et al. 2011) (Lund, Dixit et al. 2023). In some cases, coevolution promotes speciation (Sorenson, Sefc et al. 2003, Langmore, Grealy et al. 2024) and in others the evolution and maintenance of host-specific matrilines (Spottiswoode and Stevens 2011, Fossoy, Sorenson et al. 2016, Spottiswoode, Tong et al. 2022), with important genomic consequences.

Robust genomic resources for representative parasitic birds are needed to facilitate analyses of both convergence and the varied lineage-specific adaptations produced in their coevolutionary arms races with hosts (Sorenson and Payne 2002) (Ronka, Eroukhmanoff et al. 2024). Towards this end, we formed the Brood Parasitic Bird Genomes Consortium (BPBGC) and generated new genome assemblies for ten brood parasites that, combined with a recently published genome assembly for common cuckoo *Cuculus canorus* (Merondun, Marques et al. 2024) provides at least one representative for each of the seven evolutionary origins of obligate parasitism in extant birds. In addition, the genomes for eight nesting/parental bird species were assembled to provide outgroup comparisons for different parasitic lineages (Table 1). Three of the 18 new genomes reported here were produced in collaboration with the Vertebrate Genomes Project (Rhie, McCarthy et al. 2021). Female samples were chosen for all genome assemblies to include the female-specific W chromosome, which along with the mitochondrial DNA (mtDNA), is expected to be an important target of selection in some brood parasitic lineages e.g. (Spottiswoode, Stryjewski et al. 2011) (Spottiswoode and Stevens 2011, Fossoy, Sorenson et al. 2016) (Spottiswoode, Tong et al. 2022).

**Table 1.**
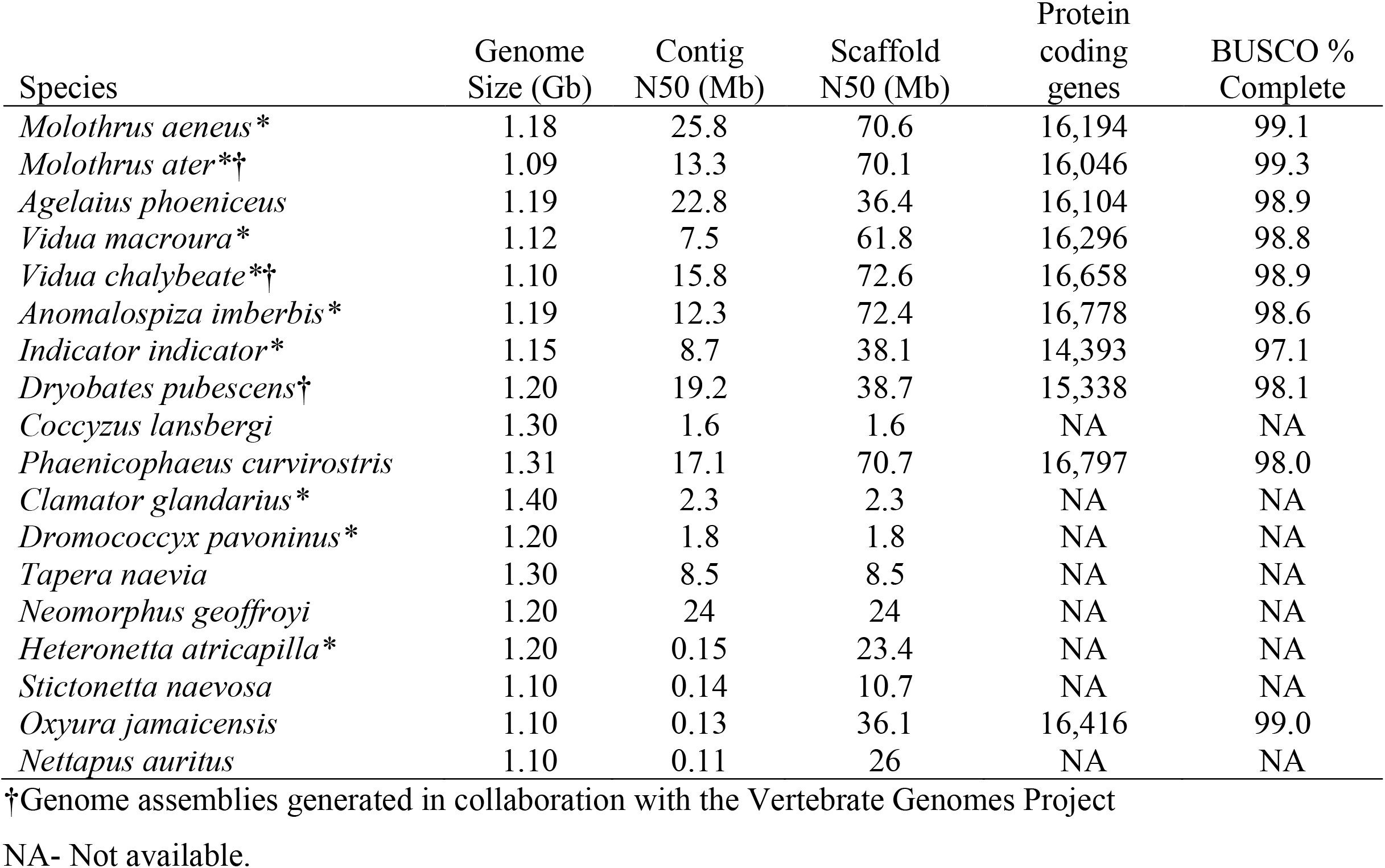
Reference genome assembly statistics for all brood parasitic* and parental outgroup assemblies generated by the BPBGC. Genomes representing the seven independent origins of obligate brood parasitism are indicated in Fig. 3, with the recently published common cuckoo genome (Merondun, Marques et al. 2024) representing one of the seven origins. For RefSeq assemblies the number of annotated protein-coding genes and completeness for BUSCO genes are provided. We report for the contig only assemblies the same N50 contig and scaffold length metrics.

| Species | Genome Size (Gb) | Contig N50 (Mb) | Scaffold N50 (Mb) | Protein coding genes | BUSCO % Complete |
| --- | --- | --- | --- | --- | --- |
| *Molothrus aeneus* * | 1.18 | 25.8 | 70.6 | 16,194 | 99.1 |
| *Molothrus ater* *† | 1.09 | 13.3 | 70.1 | 16,046 | 99.3 |
| *Agelaius phoeniceus* | 1.19 | 22.8 | 36.4 | 16,104 | 98.9 |
| *Vidua macroura* * | 1.12 | 7.5 | 61.8 | 16,296 | 98.8 |
| *Vidua chalybeata* *† | 1.10 | 15.8 | 72.6 | 16,658 | 98.9 |
| *Anomalospiza imberbis* * | 1.19 | 12.3 | 72.4 | 16,778 | 98.6 |
| *Indicator indicator* * | 1.15 | 8.7 | 38.1 | 14,393 | 97.1 |
| *Dryobates pubescens*† | 1.20 | 19.2 | 38.7 | 15,338 | 98.1 |
| *Coccyzus lansbergi* | 1.30 | 1.6 | 1.6 | NA | NA |
| *Phaenicophaeus curvirostris* | 1.31 | 17.1 | 70.7 | 16,797 | 98.0 |
| *Clamator glandarius* * | 1.40 | 2.3 | 2.3 | NA | NA |
| *Dromococcyx pavoninus* * | 1.20 | 1.8 | 1.8 | NA | NA |
| *Tapera naevia* | 1.30 | 8.5 | 8.5 | NA | NA |
| *Neomorphus geoffroyi* | 1.20 | 24 | 24 | NA | NA |
| *Heteronetta atricapilla* * | 1.20 | 0.15 | 23.4 | NA | NA |
| *Stictonetta naevosa* | 1.10 | 0.14 | 10.7 | NA | NA |
| *Oxyura jamaicensis* | 1.10 | 0.13 | 36.1 | 16,416 | 99.0 |
| *Nettapus auritus* | 1.10 | 0.11 | 26 | NA | NA |

†Genome assemblies generated in collaboration with the Vertebrate Genomes Project
NA- Not available.

## Methods

### Biological Materials

Samples of female birds were acquired from museum collections, live birds in captivity, or from materials obtained in past or current field research by us and colleagues, with the appropriate permits or MTA agreements (Supplemental Table 1).

### Sequencing

High molecular weight DNA (HMW DNA) was isolated from female muscle samples for each of the following 14 species that were designated for consensus long-read (CLR) or circular consensus sequence (CCS/HiFi) sequencing: CLR (brown-headed cowbird, *Molothrus ater;* village indigobird, *Vidua chalybeata*; greater honeyguide; *Indicator indicator* and downy woodpecker; *Dryobates pubescens*) and HiFi reads (bronzed cowbird, *Molothrus aeneus;* red-winged blackbird, *Agelaius phoeniceus;* cuckoo finch, *Anomalospiza imberbis*, pin-tailed whydah, *Vidua macroura;* great spotted cuckoo, *Clamantor glandarius*; striped cuckoo, *Tapera naevia*; pavonine cuckoo, *Dromococcyx pavoninus*; grey-capped cuckoo, *Coccyzus lansbergi*; chestnut-breasted malkoha, *Phaenicophaeus curvirostris*; and rufous-vented ground cuckoo, *Neomorphus geoffroyi*). SMRTbell libraries for sequencing were prepared according to the manufacturer’s recommended protocol (Pacific Biosciences-PacBio; Menlo Park, CA). In brief, HMW DNA was sheared to a target size of ∼15–20 kb using a Megaruptor 3 system (Diagenode), followed by DNA end-repair and barcode adaptor ligation to generate circular SMRTbell templates for sequencing. Size selection was then performed using BluePippin (Sage Science) to enrich for fragments ≥15 kb. All SMRTbell libraries were sequenced on a PacBio instrument.

Over the course of this study instrumentation upgrades occurred that included Sequel I, II, RSII, or Revio systems. The relevant sequencing mode, e.g. CLR or HiFi, was run according to the manufacturers instrument-specific protocols. Instrument use per species is specified in Supplemental Table 1. All relevant software used in this study for the purpose of genome assembly and curation, as well as any downstream genome analyses is presented in Table 2.

**Table 2.** Study software used.

| Software and any non-default options | Version or access date | Reference |
| --- | --- | --- |
| **De novo assembly** |  |  |
| Canu | 2.1 | (Koren, Walenz et al. 2017) |
| FALCON | 1.3.0 | (Biosciences 2019) |
| Flye | 2.8.1 | (Kolmogorov, Yuan et al. 2019) |
| HiFiasm | 2021-2023 | (Cheng, Concepcion et al. 2021). |
| Supernova2 | 2.1.1 | (Visendi 2022) |
| **Scaffolding and chromosome formatting** |  |  |
| Chromap | 0.2.3 | (Zhang, Song et al. 2021) |
| SALSA | 2.2 | (Ghurye, Rhie et al. 2019) |
| Juicebox | 2.3.6 | (Robinson, Turner et al. 2018) |
| Juicertools | 1.6-3.0.0 | (Durand, Shamim et al. 2016) |
| YaHS | 1.2a.1 | (Zhou, McCarthy et al. 2023) |
| Bionano Solve | 3.2.1 | (Bionano. 2019) |
| Scaff10x | 4.1.0 | (Institute 2020) |
| Longranger Align | 2.2.2 | (Genomics. 2018) |
| Agptools | 1.0 | (Ricemeyer 2025) |
| **Assembly polishing and purging** |  |  |
| Arrow | 6.0 | (Pacific Biosciences of California 2019) |
| freebayes | 1.3.1-1.3.5 | (Garrison 2012) |
| Minimap2 | 2.21 | (Li 2018) |
| PurgeHaplotigs | 1.1.3 | (Roach, Schmidt et al. 2018) |
| **Assembly QC** |  |  |
| BUSCO | 4.1.4 | (Simao, Waterhouse et al. 2015) |
| **Gene orthology and chromosome synteny** |  |  |
| OrthoFinder | 2.5.4 | (Emms and Kelly 2019) |
| Genespace | 2022 | (Lovell, Sreedasyam et al. 2022) |
| **Z chromosome evaluation** |  |  |
| BOWTIE2 | 2.5.1 | (Langmead & Salzberg, 2012) |
| mosdepth | 0.3.1 | (Pedersen & Quinlan 2018) |
| ANGSD | 0.940 | (Korneliussen et al. 2014) |
| **Phylogenetic analysis** |  |  |
| PHYLUCE | 1.7.1 | (Faircloth 2016) |
| BEAST | 2.7.7 | (Bouckaert, Vaughan et al. 2019) |

We also isolated HMW DNA from four duck species: black-headed duck (*Heteronetta atricapilla*), freckled duck (*Stictonetta naevosa)*, ruddy duck (*Oxyura jamaicensis*), and African pygmy goose (*Nettapus auritus*) to generate long DNA fragment length linked-read libraries (10x Genomics, Pleasonton, CA) for each. Library preparation was carried out with the 10x Genomics (Pleasanton, CA) Chromium^TM^ Genome Library and Gel Bead Kit v2 as described in Zhang et al. (Zhang, Zhou et al. 2019). We sequenced each linked-read library at 100bp read length to a targeted 60x depth of coverage on an Illumina NovaSeq 6000 instrument.

All sequences for each species in this study, regardless of type, have been deposited in the NCBI sequence read archive (SRA). All relevant data are retrievable by searching NCBI Bioproject accession numbers provided in Supplemental Table 1).

### Genome assembly and curation

Over the course of this project, we performed de novo genome assembly on CLR or HiFi sequence data using a variety of algorithms. Each assembly algorithm was chosen and deployed based on the type of data generated and current best practices at the time of sequence data availability and its type. We specify the assembly algorithm used for each species in Supplemental Table 1. Overall, we used one of four programs to assemble long-reads throughout the course of this study: Canu (Koren, Walenz et al. 2017), FALCON (Biosciences 2019), Flye (Kolmogorov, Yuan et al. 2019) or Hifiasm (Cheng, Concepcion et al. 2021). All primary assemblies were subjected to a version of PurgeHaplotigs (Roach, Schmidt et al. 2018) that was run on all contig assemblies to remove potential alternate haploid contigs using a custom nextflow pipeline (Warrenlab 2022).

For the nine species that were designated for chromosome scale assemblies using either CLR or HiFi reads (greater honeyguide, downy woodpecker, brown-headed cowbird, bronzed cowbird, red-winged blackbird, cuckoo finch, pin-tailed whydah, village indigobird, and chestnut-breasted malkoha) chromatin proximity interaction maps were generated for contig scaffolding. High-throughput Chromosome Conformation Capture (Hi-C) libraries were prepared using the Arima Genomics (Carlsbad, CA) or Phase Genomics (Seattle, WA) Hi-C pipelines by following the manufacturer’s standard protocols for each. All Hi-C libraries were sequenced on an Illumina Novaseq 6000 (San Diego, CA) instrument using a saturation mapping estimation of 100M paired end reads per Gb of genome size with bird genomes on average estimated to be 1.2Gb in length.

Chromosome scaffolding was performed using combinations of various tools that included Chromap (Zhang, Song et al. 2021), SALSA2 (Ghurye, Rhie et al. 2019), Juicebox (Robinson, Turner et al. 2018), Juicertools (Durand, Shamim et al. 2016), and YaHS (Zhou, McCarthy et al. 2023). We provide the combination of mapping programs used for each species in Supplemental Table 1. For all draft assemblies, including those based on CLR or HiFi read sources, base error was corrected with Arrow. An additional polishing step to correct assembly base call error for species with CLR data input was performed on greater honeyguide, downy woodpecker, and village indigobird using Illumina sequence data sourced from the same sample. Trimmomatic was used to trim adaptors, low-quality bases, and contaminants from Illumina sequences prior to BWA-MEM alignment and base call correction. Iterative polishing was completed with freebayes integrated into the VGP assembly pipeline v.1. (VGPassembly 2019) or a custom cextflow pipeline (Ricemeyer 2025). Briefly, this later pipeline aligns short reads to the assembly with minimap2 v2.21 (Li 2018), calls variants using freebayes v1.3.5 (Garrison 2012), and then edits the assembly to match the consensus from the variant calls. We ran this pipeline twice iteratively for each CLR assembly, except for the brown-headed cowbird, for which there was insufficient tissue from the same sample to generate short reads.

After polishing a sequential process was employed to build and format chromosomes by assembling the longest-range scaffolds possible for each species using Hi-C associated procedures previously described in Rhie et al. (Rhie, McCarthy et al. 2021). Briefly, Hi-C reads were mapped to the draft genome assembly using BWA-MEM, alignments were processed to generate a contact map, and resulting scaffolds iteratively ordered and oriented primarily with SALSA (Ghurye, Rhie et al. 2019), followed by manual correction of scaffolding errors the Juicebox Assembly Tools interface (Robinson, Turner et al. 2018). The pseudo-chromosomes were aligned to the assembled chromosomes of the most closesly related bird species available at that time to manually examine discrepancies and fix gross assembly errors. Agptools was used to finalize chromosome formatting (Ricemeyer 2025). For consistency and future comparative studies, all autosomes were numbered according to total size starting with the largest in accordance with the VGP guidelines (Rhie, McCarthy et al. 2021). As a result, the numerical assignment of chromosomes will not always reflect homology across species.

The genomes of the duck clade including the black-headed duck, freckled duck, ruddy duck, and African pygmy goose were assembled with Supernova v2.1.1 (Visendi 2022) following the methods outlined in Zhang et al. (Zhang, Zhou et al. 2019). Initial chromosomal assignments were determined by aligning assembly scaffolds from each species to the reference genomes of domestic chicken (*Gallus domesticus*, GCA_000002315.5) and the mallard (*Anas platyrhynchos*, GCA_003850225.1). To improve chromosomal accuracy, we conducted iterative pairwise alignments among the four duck genomes, allowing each to inform the structural organization of the others. Final chromosome numbering was assigned based on synteny with the chicken genome.

All chromosome assemblies were scanned for repetitive sequences using WindowMasker (Morgulis, Gertz et al. 2006), enabling the resulting output to be utilized for future genome masking steps in downstream analyses. Assembly completeness was evaluated using BUSCO v4.1.4 with the aves_odb10 lineage dataset to assess conserved gene content coverage (Simao, Waterhouse et al. 2015). Additionally, corrections made to coding sequence (CDS) predictions using the Splign alignment algorithm (Kapustin 2008) were used as an independent measure of consensus sequence accuracy, with the number of CDS corrections reflecting potential base-calling errors.

### Gene Annotation

Gene annotations for all chromosome-scale assemblies were generated using the NCBI Eukaryotic Genome Annotation Pipeline, as described in Rhie et al. (Rhie, McCarthy et al. 2021). RNA-seq datasets from diverse tissue types, publicly available in the NCBI SRA for each species, were integrated into the annotation process to inform and support transcript model predictions. As part of this process, protein alignment to the genome was used to generate preliminary predicted models, and to correct as needed the final models with compensating insertions or deletions to correct for spurious indels in the genome. These errors were tallied as an additional comparative metric of annotation quality.

### Gene orthology

To compare gene orthology across the avian genomes in this study, we ran OrthoFinder version 2.5.4 using the RefSeq protein-coding gene predictions for each species as input (Emms and Kelly 2019). To mitigate the effect of multiple transcripts per gene, we used only primary transcripts (the longest version of protein) only, per OrthoFinder recommendations (Emms and Kelly 2019).

### Whole genome interspecies synteny analysis

To explore shared gene organization across parasitic species and parental outgroups, we analyzed syntenic orthology among species using the Genespace R package (Lovell, Sreedasyam et al. 2022). The parse annotation function in Genespace was used to extract a gene Browser Extensible Data (BED) file from the General Feature Format (GFF) files of each species found in the NCBI Reference Sequence Database. After ensuring proper formatting and matching headers between protein and gene BED files, we initiated Genespace as described in Lovell et al (Lovell, Sreedasyam et al. 2022). Pairwise comparisons within clades revealed that, in some cases, entire chromosomes appeared to be inverted end-to-end. However, these apparent reversals were attributed to species-specific differences in chromosomal orientation in our assemblies, rather than true biological inversions. To ensure more accurate representation of syntenic relationships across species, such chromosomes were manually reoriented for the purpose of visualization.

### Z chromosome evaluation in downy woodpecker

To better understand the unusually long Z-chromosome in down woodpecker, we mapped short-read sequencing data to the reference genome using BOWTIE2 v. 2.5.1 (Langmead & Salzburg, 2012) and then used mosdepth v. 0.3.1 (Pedersen & Quinlan 2018) to calculate sequencing depth in 10 kbp windows for two male and four female downy woodpeckers. We used ANGSD (Korneliussen et al. 2014) to calculate individual heterozygosity in 50 kbp windows for these same six individuals.

### Phylogenetic Analysis

Sequence data from ultraconserved elements (UCEs) provide a powerful tool for estimating a time-calibrated phylogeny (time tree) of avian species and offer an opportunity to investigate the evolutionary origins of obligate brood parasitism. We used UCEs to estimate a time tree for 52 species comprising representatives of the seven lineages of obligate brood parasites, closely related nesting relatives, and other selected lineages in the avian tree of life (Supplemtnal Table 2). UCEs were extracted from whole genome assemblies using PHYLUCE v1.7.1 (Faircloth 2016) and a probe set designed to target ∼5,000 loci (Faircloth, McCormack et al. 2012). We extracted 1,000 bp of the flanking regions for each locus, and sequences for individual loci were aligned and edge trimmed. Loci with < 47 taxa represented were filtered out, resulting in 4,491 retained UCE loci that were concatenated and used for phylogenetic inference in BEAST v2.7.7 (Bouckaert, Vaughan et al. 2019). Based on results from a recent phylogenomic study of birds, we constrained the analysis with several monophyletic priors, including monophyly of most families and key higher-level relationships (Stiller, Feng et al. 2024). We also adopted six fossil-based node age calibrations from this recent study (Stiller, Feng et al. 2024), applying them as log-normal distribution priors to estimate branch lengths in units of millions of years. We initiated 10 independent runs with different starting points in BEAST, each with the GTR+I+G model of sequence evolution, a birth-death coalescent prior, and sampling every 1,000 generations. Runs were stopped after 5M generations, and inspection of posteriors found that six of the runs reached stationarity at similar likelihood scores. After removing the appropriate burn-in for each run, results were combined across these six runs to form a posterior sample of 10,000 trees that were used to estimate a maximum clade credibility time tree.

## Results and Discussion

### Chromosome scale assemblies of long-read sequenced avian genomes

Among the 18 avian genome assemblies we generated over a four-year span, ten were assembled to the chromosome level using only CLR or HiFi input sequences; these include both brood parasitic and non-parasitic outgroup species (Table 1; Supplemental Table 1). Female samples were chosen for all species to allow analyses of the W-chromosome. Considering all sequence technologies employed, we achieved a total depth of sequence coverage ranging from 30x to 254x irrespective of sequence type (Supplemental Table 1). The total assembly sizes exhibited a narrow range of 1.1 to 1.3 Gb with N50 contig and scaffold length, respectively, from 7.49 to 25.8 Mb and 38.1 to 72.6 Mb, respectively (Table 1). Improvements in assembly contiguity were achieved as sequencing capacity and accuracy improved along with advancements in de novo assembly algorithms, e.g. PacBio CLR compared to HiFi inputs (Table 1; Supplemental Table 1). This is illustrated in our most recently completed assembly, the bronzed cowbird, which has the highest metrics of contiguity (Table 1). Despite the improvements in overall assembly quality between earlier and later assembled species, we judge all the chromosome scale references to be highly contiguous and of sufficient accuracy for various comparative genomic analyses.

### Contig-level assemblies for additional cuckoo species

Contig level assemblies were generated from PacBio/HiFi data for greater representation of cuckoo species, including great spotted cuckoo, grey-capped cuckoo, pavonine cuckoo, striped cuckoo, and rufous-vented ground cuckoo. A HiFi sequence coverage depth of 30x was achieved for all except the great spotted cuckoo at 19x (Supplemental Table 1). Total assembled genome sizes range from 1.2 to 1.39 Gb with contig N50 length of 1.6 to 8.5 Mb (Supplemental Table 3). For these cuckoo species, no contig scaffolding or chromosome-scale formatting was completed. Nonetheless, these contig-only assemblies offer researchers broader access to the cuckoo clade, including two additional independent origins of obligate brood parasitism, and valuable resources for gene-level analyses and population genetic studies.

### Chromosome-scale assemblies of linked-read sequenced duck genomes

Chromosome-scale assemblies were also generated for four duck species: the brood parasitic black-headed duck, and representatives of three closely related, non-parasitic genera, the African pygmy goose, ruddy duck, and freckled duck. However, all these assemblies were generated with long range fragment sequences from the 10X pipeline as described previously (Zhang, Zhou et al. 2019). Assembly contiguity statistics were as expected for this approach with ranges of N50 contig and scaffold lengths being 105 to 151kb and 11 to 36Mb, respectively (Supplemental Table 3). Despite the high-level scaffolding contiguity achieved in these assemblies the total contig numbers remain high due to the short-read assembler input (Supplemental Table 3).

### Autosomal synteny

Based on an analysis framework of conserved gene order and single-copy orthologs use, we show a substantial level of shared chromosomal synteny within selected phylogenetic clades, but notable species pairwise species differences as well, providing first approximations of how chromosome order and orientation changed over varied spans of avian evolutionary divergence (Fig. 1). The autosomal alignments for piciforms, comprising downy woodpecker and greater honeyguide overall displayed the most extensive number of rearrangements (“Clade 3” in Fig. 1). For example, the downy woodpecker chromosome 1 corresponds to greater honeyguide chromosomes 8 and 15. Because the greater honeyguide assembly was produced from lower-quality CLR data (Supplemental Table 1), we cannot rule out an assembly artefact as an explanation for this result, although we note that woodpecker genomes are reported to have an unusually high number of rearrangements for birds (de Oliveira, Kretschmer et al. 2017, Hruska and Manthey 2021, Forest, Achaz et al. 2024). In contrast, most other pairwise alignments by clade display relatively high levels of chromosomal synteny (Fig. 1), interspersed with some inversions between closely related species. We note that additional orthogonal evidence will be needed to confidently establish whether the breaks in synteny that we observed in this study represent assembly error or natural structural variation (Fig. 1).

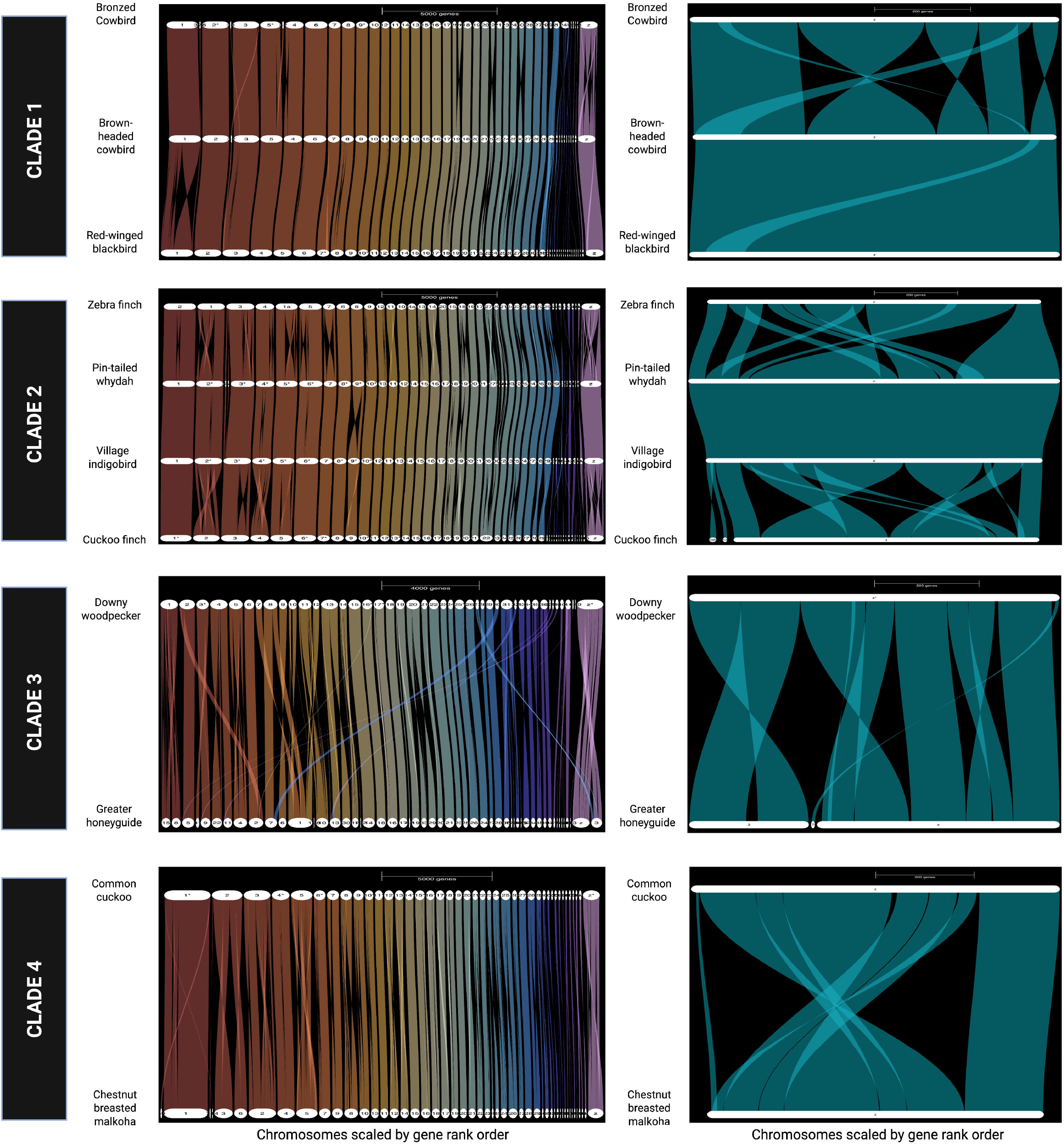

### Sex chromosomes assembly and synteny

Given evidence for maternal inheritance of egg phenotypes (Spottiswoode, Stryjewski et al. 2011) phenotypes (Spottiswoode and Stevens 2011, Fossoy, Sorenson et al. 2016) (Spottiswoode, Tong et al. 2022) (Merondun, Marques et al. 2024), we evaluated the Z and W chromosomes for possible sources of genetic variation relevant to host-specific adaptation (Fig. 1). The assembled sizes of Z chromosomes ranged from 70 to 84 Mb (Supplemental Table 4), with one notable exception: the downy woodpecker, which was 140 Mb. The downy woodpecker Z chromosome largely aligns with the honeyguide Z chromosome, but a substantial portion of its sequence aligns with chromosome 3, suggesting a possible autosomal translocation (Fig. 1). Autosomal translocations or fusions involving an autosome, and the sex chromosomes, yielding “neo-sex chromosomes”, have been reported in songbirds (Sigeman, Strandh et al. 2021) (Sigeman, Zhang et al. 2022) and parrots (Huang, De et al. 2022), though these events remain rare among avian species. Our proximity mapping analysis strongly supports a translocation or fusion of a portion of an autosome to the sex chromosomes in the history of woodpeckers, the precise timing of which is uncertain without comparative data from other species (Supplemental Figure 1). The assembled size of the downy woodpecker Z chromosome in this study is consistent with woodpecker karyotypes, which include an enlarged Z chromosome (Shields et al. 1982). The mapping of whole genome resequencing data for male and female downy woodpeckers also provides support for a Z-autosome fusion (Supplemental Figures 2–3). Relative sequencing depth for males is consistent across the full length of the Z-chromosome assembly, whereas relative depth for females increases from 0.5 to 1 at the putative junction between the ancestral Z-chromosome and former autosome (Supplemental Figure 2). This suggests that the autosomal segment fused to both the Z and W chromosomes and that divergence between the neo-Z and neo-W for the formerly autosomal segment remains sufficiently low that both W-linked and Z-linked sequences in females align to the Z-chromosome. This also assumes that the formerly autosomal portion of the neo-W is not in the genome assembly, such that females appear to be diploid for this region. To the extent that recombination suppression has evolved, divergence between the neo-Z and neo-W chromosomes should generate higher estimates of apparent heterozygosity in females than in males, but we found no evidence to support this (Supplemental Figure 3), suggesting that the formerly autosomal portions of the neo-Z and neo-W remain pseudoautosomal in downy woodpeckers. Our analysis of other clades detected multiple Z chromosome inversions of potential interest (Fig. 1). As noted earlier for autosomes, apparent Z chromosomes structural rearrangements will also require further experimental validation prior to interpreting their evolutionary significance.

W-chromosome assemblies ranged in size from 3 to 12 Mb, excluding the greater honeyguide W, underscoring the persistent difficulty of fully resolving sex-limited chromosomes (W or Y), even with high-accuracy long-read sequencing. Avian W chromosomes, like mammal Ys, are enriched in repeat content, structure, and degeneration (Huang, Xu et al. 2023) (Benham, Cicero et al. 2024), and both the Y and W represent classic examples of sex chromosome specialization and degenerative evolution, leading to compact, repeat-rich, gene-poor chromosomes shaped by similar evolutionary pressures (Bellott, Skaletsky et al. 2017). Even the chicken W chromosome, the most extensively studied in birds remains incomplete despite the application of multiple long-read technologies (Huang, Xu et al. 2023). Notably, the greater honeyguide W chromosome was particularly fragmented, with the longest W-linked contig spanning only 0.9 Mb (Supplemental Table 4). This likely reflects both high assembly fragmentation due to the more error-prone CLR data for this genome (Supplemental Table 4), and as a result lower corroboration of mapping evidence. We mostly failed to align greater honeyguide scaffolds at sufficient nucleotide similarity thresholds to the downy woodpecker W assembly, the most closely available genome (Supplemental Table 4). For all avian W chromosomes it will be necessary to implement recent technological developments that leverage complementary long-read platforms, such as PacBio HiFi and Oxford Nanopore ultra-long reads, alongside the use of evolving algorithms like Verkko (Rautiainen, Nurk et al. 2023). This approach has resulted in generating gap-free assemblies of structurally complex sex chromosomes such as the complete human Y chromosome (Rhie, Nurk et al. 2023), a result relevant for future research on the avian W.

### Gene representation and annotation of chromosome assemblies

Benchmark Universal Single Copy Orthologue (BUSCO) analysis (Simao, Waterhouse et al. 2015) revealed all chromosome-level assemblies to have a completeness score >97% (Supplemental Table 5). Importantly, the percentage of missing genes exceeded 2.6% for the greater honeyguide, whereas all others were below 1.6% (Supplemental Table 5). The number of predicted protein-coding genes using the NCBI RefSeq Annotation pipeline (O’Leary, Wright et al. 2016) per species ranged from 14,393 to 16,797 (Table 1). Estimates of noncoding gene classifications varied significantly across species (Supplemental Table 6), primarily reflecting the availability of species-specific RNAseq data. The number of long non-coding RNAs (lncRNAs) ranged from 45 to 10,129 (Supplemental Table 6). Notably, the greater honeyguide genome is predicted to have only 45 lncRNAs, likely due to its limited RNA-seq data, which was derived from one blood sample with 99 million reads. In contrast, the cuckoo finch genome was annotated using 10 billion RNAseq reads with greater tissue diversity, resulting in 10,129 predicted lncRNAs. This example highlights the need for more extensive RNA sequencing of diverse tissue sources to further improve genome annotations, both for the species characterized here and more generally.

### Gene orthology

Understanding the genetic underpinnings of adaptations linked to the evolution of brood parasitism and the coevolution of brood parasites with host species, depends on a robust gene orthology framework to enable precise investigations of genotype-to-phenotype relationships. Orthologs are genes that diverged at a speciation event, whereas paralogs are genes that arise from gene duplication events. Using OrthoFinder, we inferred orthogroups, identified species-specific orthogroups, analyzed gene orthogroup multiplicity, and estimated gene duplications at various evolutionary nodes (Fig. 2A-G). The fraction of genes in orthogroups was similar across species (Fig. 2B), although the number of species-specific orthogroups and shared orthogroups were not. The common cuckoo and chestnut-breasted malkoha exhibited the highest number of species-specific orthogroups, defined as sets of genes descended from a single ancestral gene in their last common ancestor (Fig. 2C). Meanwhile, the greater honeyguide appears to have the smallest number of shared orthogroups compared with other species (Fig. 2D). Overall, when using zebra finch as a reference, we observe a high number of shared one-to-one orthologs across all species (Fig. 2E). Among the six parasitic species compared the greater honeyguide followed by cuckoo finch had the greatest number of species-specific gene duplications (Fig. 2F). The smaller number of shared orthogroups and higher number of gene duplications in greater honeyguide is likely due to misannotated genes and genes with split annotations. The number of duplications on each branch of the species tree that are retained in all descendant species was highest for the passeriform clade, comprising cowbirds, village indigobird, and cuckoo finch (Fig. 2G). Overall, our results are in line with other studies showing that greater evolutionary divergence between species will correlates with a reduced number of one-to-one gene orthologs (Emms and Kelly 2019).

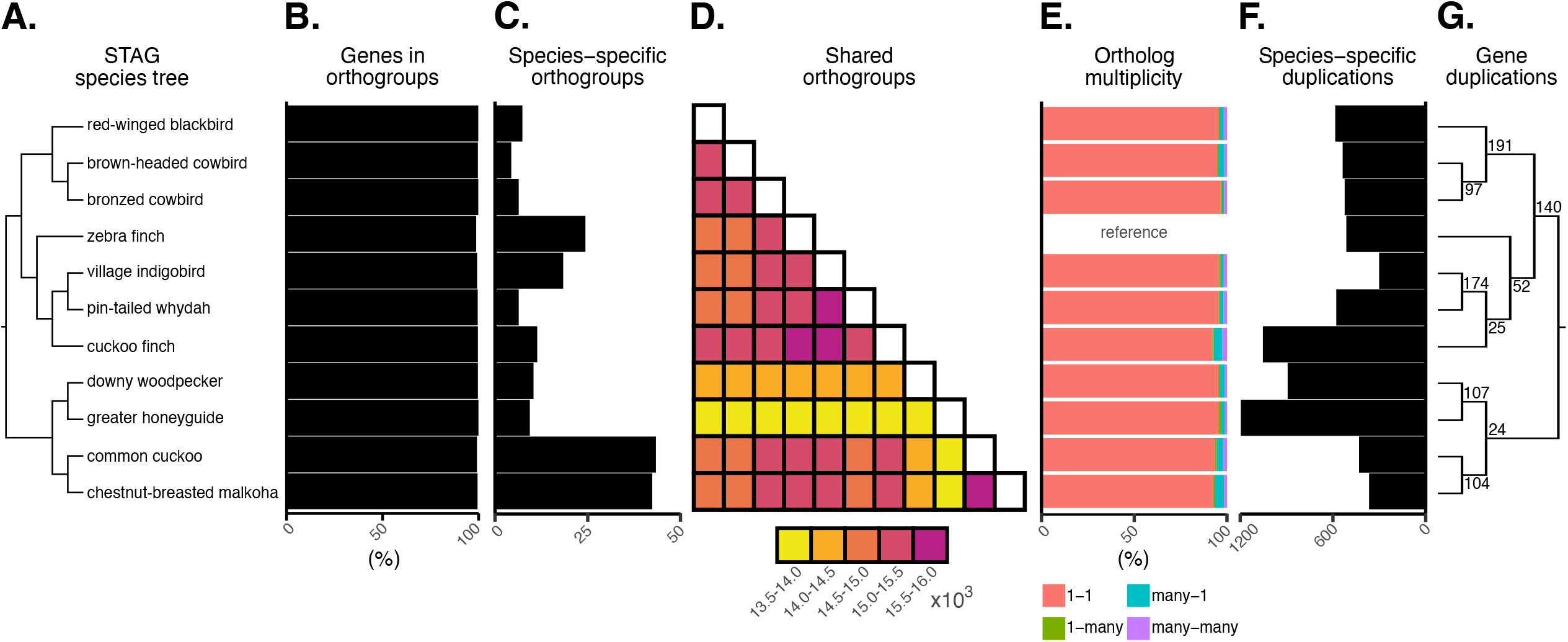

### Evolutionary origins of brood parasitism in phylogenetic context

Our phylogenomic analysis using concatenated UCE loci resulted in six independent runs that converged in similar parameter space; these runs were combined in a posterior sample of 10,000 trees. Family and higher-level relationships were constrained, guided by the results of Stiller et al. (Stiller, Feng et al. 2024). Our improved sampling of brood parasites provides a single, robust analysis for comparing the timing of the seven independent origins of obligate brood parasitic behavior (Fig. 3). The most recent evolution of brood parasitism was in cowbirds (*Molothrus*), with point estimates of minimum and maximum ages at 1.6 and 4.5 mya, respectively, with the minimum estimate taken from the common ancestor of brown-headed and bronze cowbird and the maximum estimate taken from the common ancestor of cowbirds and the most closely related parental species (i.e., the two ends of the ancestral cowbird lineage). Applying the same logic to estimates for the origin of brood parasitism in parasitic finches (*Anomalospiza*-*Vidua*; 13.7-16.8 mya), honeyguides (*Indicator*; 10.4-29.2 mya), and the *Dromococcyx*-*Tapera* clade of cuckoos (15.3-34.5 mya) indicates comparatively older origins of parasitism. With sampling of only a single species, minimum ages could not be estimated for the black-headed duck (*Heteronetta*; max=14.3 mya), *Calamator* cuckoos (max=17.8 mya), or the large clade of parasitic cuckoos that includes the common cuckoo (*Cuculus canorus*; max=30.5 mya). This topology with branch lengths provides an approximate timeframe for future comparative analyses of parasitic birds.

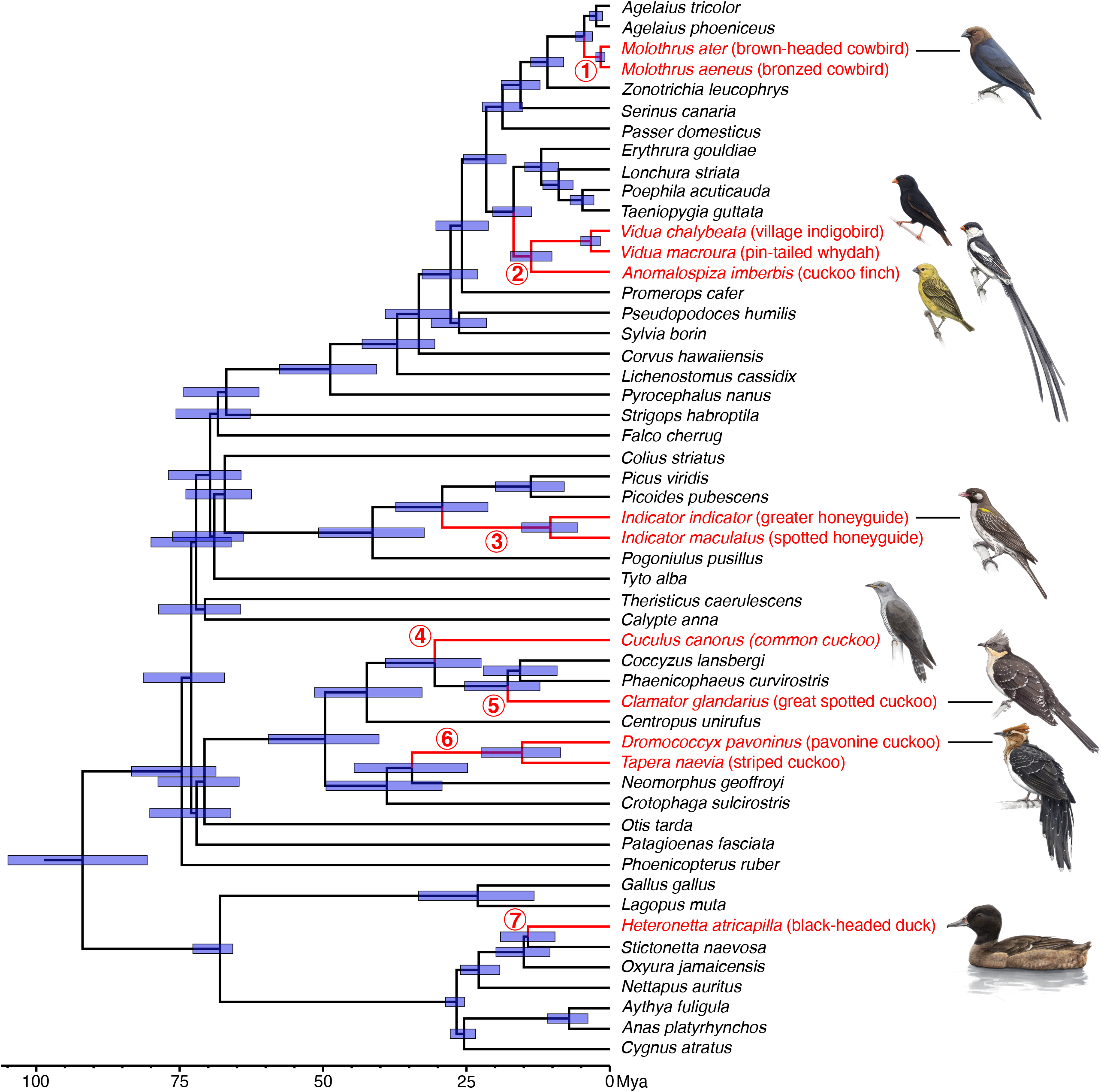

Avian brood parasitism, defined by the deceptively simple act of laying an egg in the nest of an appropriate host species, has figured prominently in evolutionary biology, providing a fascinating diversity of coevolved adaptations in interacting hosts and parasites. This research has largely been behavioral and ecological with few tests of the genetic basis of the unique adaptations of brood parasites, including their loss of parental care behaviors. This consideration figured prominently in the BPBGC’s development of the genomic resources described here. Our broader aim was to fill important taxonomic gaps in the availability of reference genomes to facilitate genomic analyses this rare but widespread reproductive strategy. In doing so, we highlight key genomic features, including chromosomal synteny and gene orthology, and provide a phylogenetic timescale for the evolution of parasitic behavior in each clade. By reporting measures of assembly quality, focusing on contiguity, synteny, and gene representation evaluations, we encourage users to select assemblies described herein based on their specific computational objectives. These genomic resources will enable more detailed testing of diverse hypotheses about the origins of brood parasitism and adaptations of parasitic birds.

## Acknowledgements

For their assistance in collecting or providing the samples needed for genome assembly and annotation, we thank Eric Schuppe, Matt Louder, Louisiana State University Museum of Natural Science, Museum of Southwestern Biology, Texas Cooperative Wildlife Collection, Beckman Center for Conservation Research at the San Diego Zoo, Sylvan Heights Bird Park, Museum of Comparative Zoology (Harvard), and the many additional individuals and institutions that supported or assisted in past fieldwork and who contributed samples to museum collections or assisted directly or indirectly in other ways. We thank Maggs X for recommendations on the parameterization of Orthofinder. We thank the members of the Hudson Alpha Institute sequencing core, especially Jane Grimwood, for long read sequencing data generation. Finally, we thank various members of the VGP consortium including Giulio Formenti, Olivier Fedrigo and Bonhwang Koo for assistance during the sequencing and assembly of the brown-headed cowbird, village indigobird, and downy woodpecker. We thank Melissah Rowe, Judith Risse, Simon Grifith and Daniel Hooper for early access to a *Poephila acuticauda* genome assembly. The work of DH, FTN and PM was supported by the National Center for Biotechnology Information of the National Library of Medicine (NLM), National Institutes of Health.

## Data availability

We have deposited the primary data underlying these analyses as follows: chromosome-level or contig-only assemblies were deposited at Genbank under various accession numbers summarized in Supplemental Table 1. All raw sequence data were deposited in the NIH’s sequence read archive under the appropriate species Bioproject identification number (Supplemental Table 1). The WGS data used to evaluate Downy woodpecker Z chromosome is found in GenBank (SRR33614311–20).

