## Supplemental Figures for "Genomic resources for comparative analyses of obligate avian brood parasitism"

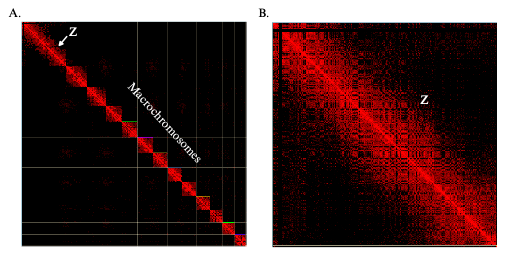

**Supplementary Figure 1**. Contact proximity maps of the Downy woodpecker chromosomes. A. The HiC heat map of contact points for the Downy woodpecker Z chromosome and largest macrochromosomes. B. The HiC contact map for the primary Z chrosomosome. Visualization of all HiC contact maps were generated with PretextSnapshot. Omni-C contact maps translate the proximity of genomic regions in 3D space to contiguous linear organization. Each cell in the contact map corresponds to sequencing data supporting the linkage (or join) between two such regions.

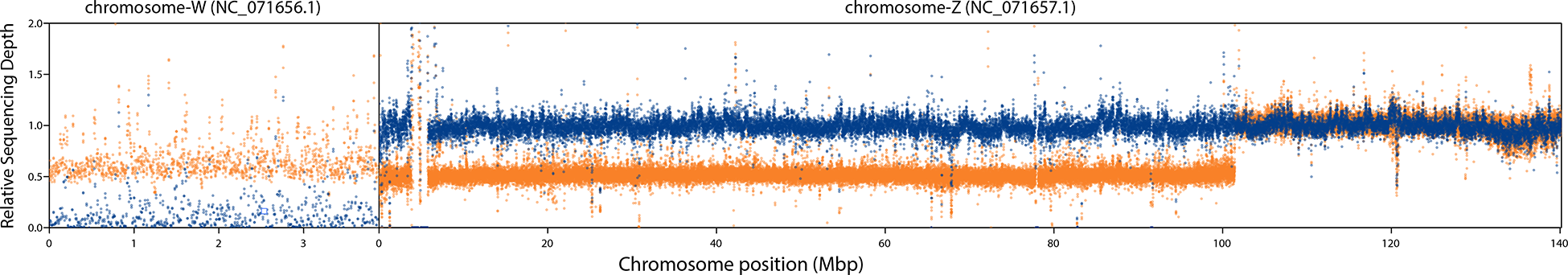

**Supplementary Figure 2.** Relative sequencing depth for two male (blue) and four female (orange) downy woodpeckers across the W- and Z-chromosome assemblies. Sequencing depth relative to the autosomal median for each sample is plotted for 10 kbp windows. As expected, males have low depth for the W-chromosome, whereas females have approximately half the autosomal depth. Also as expected, across the first ~104 Mbp of the Z-chromosome, males have twice the sequencing depth as females. For the remaining ~36 Mbp of the Z-chromosome, which shows homology to one of the larger autosomes in other piciforms, males and females have comparable sequencing depth, suggesting that the 36 Mbp region was also transposed to the W-chromosome but is not included in the genome assembly.

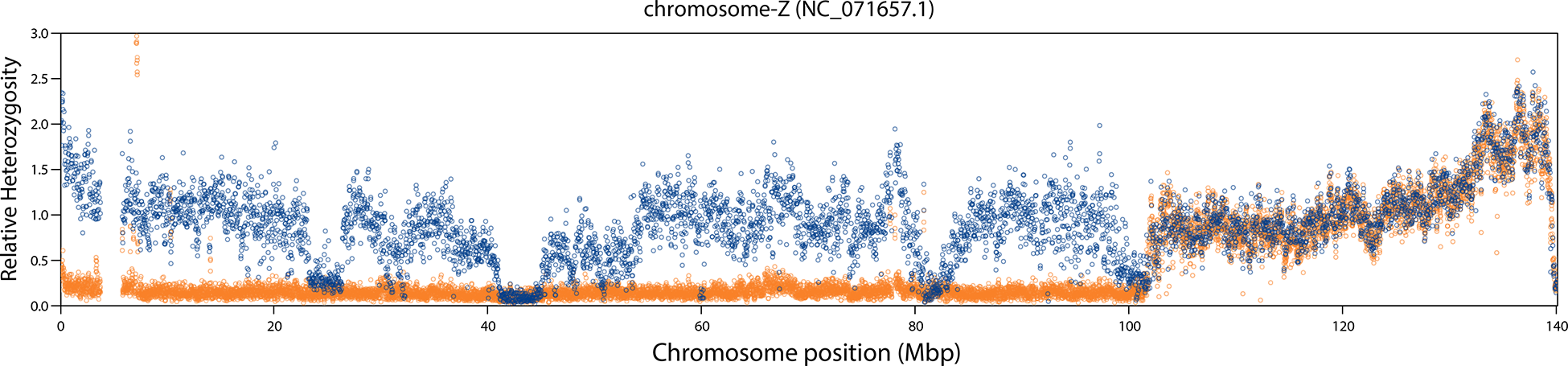

**Supplementary Figure 3**. Comparison of heterozygosity for two male (blue) and four female (orange) downy woodpeckers across the Z-chromosome assembly. Heterozygosity relative to the autosomal average for each sample is plotted for 50 kbp windows. As expected, male heterozygosity is comparable to autosomal values for most of the chromosome, whereas female heterozygosity is close to zero for the first ~104 Mbp (these values should be zero but are presumably somewhat greater than zero due to some W-chromosome sequences and/or autosomal sequences from repetitive regions aligning to the Z-chromosome). If the remaining ~36 Mbp represents a recent transposition to both the Z- and W-chromosomes and the W-linked copy of this region is not in the genome assembly, then both Z- and W-linked reads are expected to align to the Z-chromosome assembly in females. However, if the Z- and W-linked copies have begun to diverge due to cessation of recombination, then we would predict that females would have higher apparent heterozygosity in this region. The data do not support this prediction, suggesting that this portion of the Z- (and presumably W-) chromosome continues to freely recombine and is in effect pseudoautosomal. Note that the genes typically found in the small pseudoautosomal region (PAR) in other avian species map to an unplaced contig (NW_026530690.1) in the downy woodpecker assembly.
